# Transgene-mediated skeletal phenotypic variation in zebrafish

**DOI:** 10.1101/792929

**Authors:** Charles B. Kimmel, Alexander L. Wind, Whitney Oliva, Samuel D. Ahlquist, Charline Walker, John Dowd, Bernardo Blanco-Sánchez, Tom A. Titus, Peter Batzel, John H. Postlethwait, James T. Nichols

**Author notes:** **Correspondence** James T. Nichols, Craniofacial Biology, Mail Stop 8120, 12801 E. 17th Avenue, Aurora, CO 80045.

## Abstract

When considering relationships between genotype and phenotype we frequently ignore the fact that the genome of a typical animal, notably including that of a fish and a human, harbors a huge amount of foreign DNA. Some of it, including the DNA of “autonomous” transposable elements, can spontaneously mobilize to occupy new chromosomal sites and take on new functions, presenting a challenge to the host organism and also possibly introducing new fuel for evolutionary change. Transposable elements are useful for introducing transgenes, integrating them into host genomes with high efficiency. Transgenesis has become very widespread in biological research, and in our society at large. This year the governments of both Canada and the United States have approved the first use of ‘genetically engineered’ animals in food production, Atlantic salmon, *Salmo salar*. With the recent advent of amazing gene-editing technology, there is no doubt that the transgene industry will grow explosively in the coming years. The biology of transgenes needs to be included in our understanding of the genome. It is in this spirit that we have investigated an unexpected and novel phenotypic effect of the chromosomally integrated transgene *fli1a-F-hsp70l:Gal4VP16*. We examine larval *fras1* mutant zebrafish (*Danio rerio*). Gal4VP16 is a potent transcriptional activator, and already well known for toxicity and mediating unusual transcriptional effects. In the presence of the transgene, phenotypes in the neural crest-derived craniofacial skeleton, notably fusions and shape changes associated with loss of function *fras1* mutations, are made more severe, as we quantify by scoring phenotypic penetrance, the fraction of mutants expressing the trait. A very interesting feature is that the enhancements are highly specific for *fras1* mutant phenotypes – occurring in the apparent absence of more wide-spread changes. Except for the features due to the *fras1* mutation, the transgene-bearing larvae appear generally healthy and to be developing normally. The transgene behaves as a genetic partial dominant: A single copy is sufficient for the enhancements, yet, for some traits, two copies may exert a stronger effect. We made new strains bearing independent insertions of the *fli1a-F-hsp70l:Gal4VP16* transgene in new locations in the genome, and observed increased severities of the same phenotypes as observed for the original insertion. This finding suggests that sequences within the transgene, e.g. Gal4VP16, are responsible for the enhancements, rather than effect on neighboring host sequences (such as an insertional mutation). The specificity, and biological action underlying the traits, are subjects of considerable interest for further investigation, as we discuss. Our findings show that work with transgenes needs to be undertaken with caution and attention to detail.

## 1 INTRODUCTION

### 1.1 Objective

Understanding phenotypic variation presents a principal challenge at every level of biological organization (Hallgrimsson & Hall, 2005) – from the global biodiversity crisis on the one hand (https://www.nature.com/articles/d41586-019-01448-4) to within-tissue cellular competition on the other (Meyer *et al*., 2014). Here we address the considerable skeletal variation associated with a zebrafish model of a human disease, Fraser syndrome. We show that mutant phenotypes in the facial skeleton can be potentiated, made more severe, by a transgene, *fli1a-F-hsp70l:Gal4VP16*, residing in the genetic background. We pursue this work to provide deeper understanding of the mapping between genotype and phenotype, and the function of genotype– phenotype mapping in the issue of variation.

### 1.2 Background: Fraser mutants have abnormal fusions between facial skeletal elements

Fraser Syndrome is a rare, usually deadly genetically-based disease affecting tissues that include, among other organs, the eyes, digits, kidneys, and in particular, for our study, the ears. In each of these affected organs the disease appears to result from disruption of epithelial-mesenchymal interaction (Smyth and Scambler, 2005) critically dependent on basically the same extracellular protein complex, the Fraser protein complex. The Fraser protein complex normally is a component of the basal lamina underlying epithelia in a number of body organs, and functions in intercellular signaling and/or adhesion. The molecular core of the complex is made from three giant proteins, each of which is required for its function such that loss of any of them causes Fraser Syndrome. The first of these proteins to be discovered is encoded by the gene *FRAS1* (Slovotinek *et al*., 2006), and we study the zebrafish ortholog of this gene, *fras1*, in our animal model of the human disorder.

Among the malformations of the human disease are abnormal fusions among the auditory ossicles – the three cartilage replacement bones of the middle ear normally connected by joints – that serve to amplify acoustic signals. Accordingly, the fusions can cause conductive hearing loss. Whereas fishes have no middle ear, skeletal elements homologous to the human auditory ossicles are present and form jaw and jaw support structures. As is directly comparable to the human condition, we see abnormal shapes and fusions among these elements (cartilaginous in young larvae) stemming from loss of function mutations of *fras1* (Figure 1). The defects include a prominent ectopic fusion between the symplectic and ceratohyal cartilages derived from the second pharyngeal arch (sy and ch in Figure 1), and the palatoquadrate and Meckel’s cartilages of the first pharyngeal arch (pq, M), this fusion occurring where the Meckels’-palatoquadrate joint (jaw joint) would normally be present. The severities of these phenotypes vary both in penetrance – the fraction of mutants expressing a mutant phenotype, and expressivity – the degree of phenotypic abnormality of a given element in individual mutants. For example, in two largely independent studies, Talbot *et al*. (2012; 2016) determined overall penetrance of the Meckels’-palatoquadrate fusion in *fras1^te262^* mutants as 19% and 57%, representing dramatic phenotypic diversity in different clutches of homozygotes bearing the same *fras1* mutant allele. As an example of fine-level expressivity differences, Figure 2 shows the symplectic-ceratohyal phenotypes in *fras1^te262^* mutants, as compared with the wild-type (WT) condition (Figure 2A). In panels B and D, we see complete sy-ch fusions: At the fusion site, cells from both elements are thoroughly interdigitated. A difference between the two examples is that in B, a short region of unfused symplectic cartilage extends past the fusion site (to the left in the image), but such an unfused extension is missing in D. In another mutant (Figure 2C) the symplectic and ceratohyal are quite closely opposed, but not fully fused; a thin light-staining stripe of tissue extends between them. We interpret the full fusions as being the more severe disruptions (higher expressivity).

**FIGURE 1.**
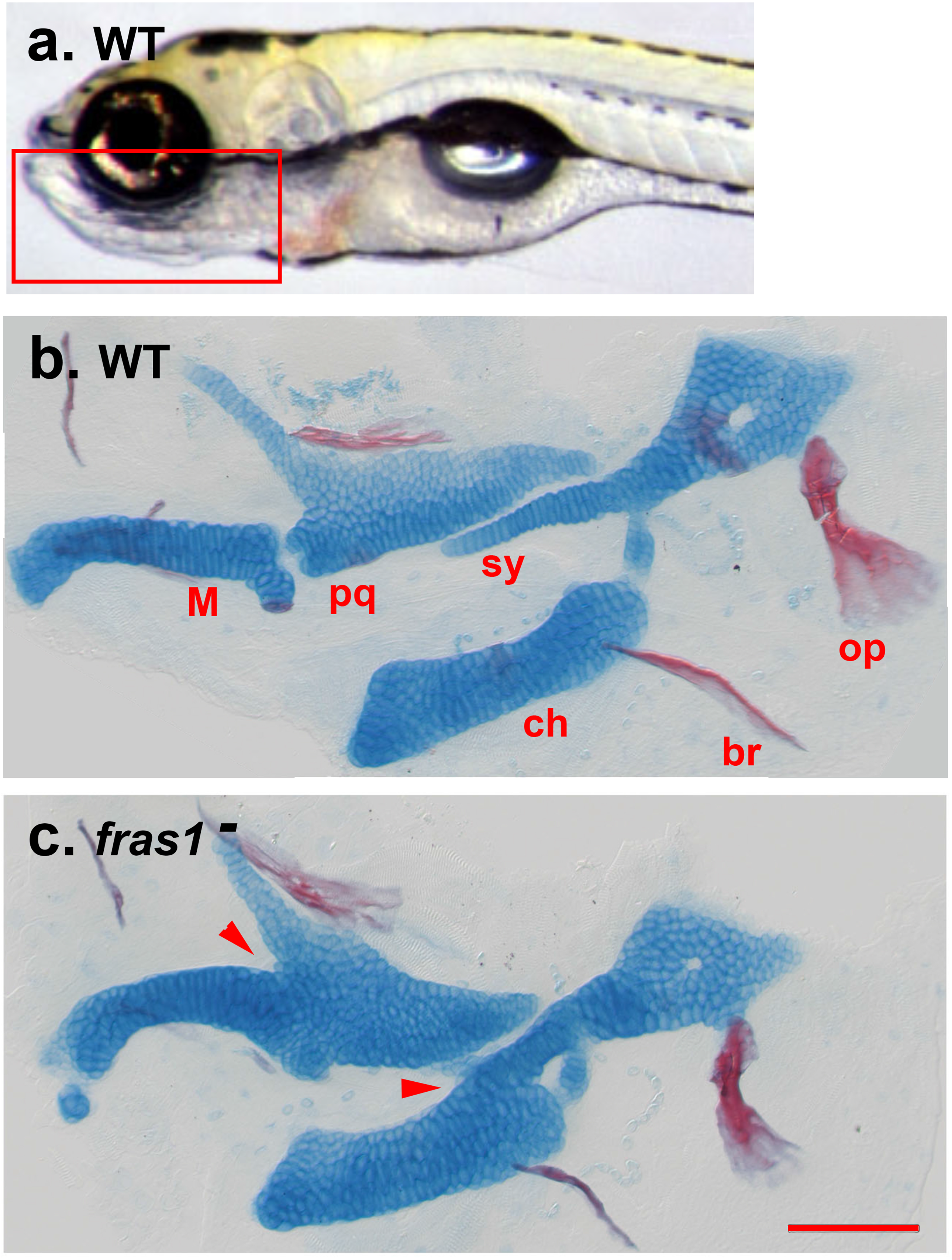
Skeletal elements in young wild type and fras1 mutant zebrafish larvae. (a) Live larva, the rectangle indicates the region of the ventral head that contains the skeletal elements of interest. Left side view, anterior to the left. (b, c) Flat mounts stained with Alcian Blue and Alizarin Red (for cartilage and bone, respectively), same orientation as A. Most of the cartilage at this stage is replaced with cartilage-replacement bone in the adult. The cartilages of interest in this study are labeled, br: branchiostegal ray, ch: ceratohyal, M: Meckel’s, pq: palatoquadrate, sy: symplectic region of the hyosymplectic. Two cartilage fusions are visible in c and indicated by arrowheads, the M-pq, and the sy-ch. The opercle (op) is a dermal bone, note its modestly reduced size in c. Abbreviations: WT: wild type, *fras1*^−^: *fras1* mutant. Scale bar, for b and c: 100µm.

**FIGURE 2.**
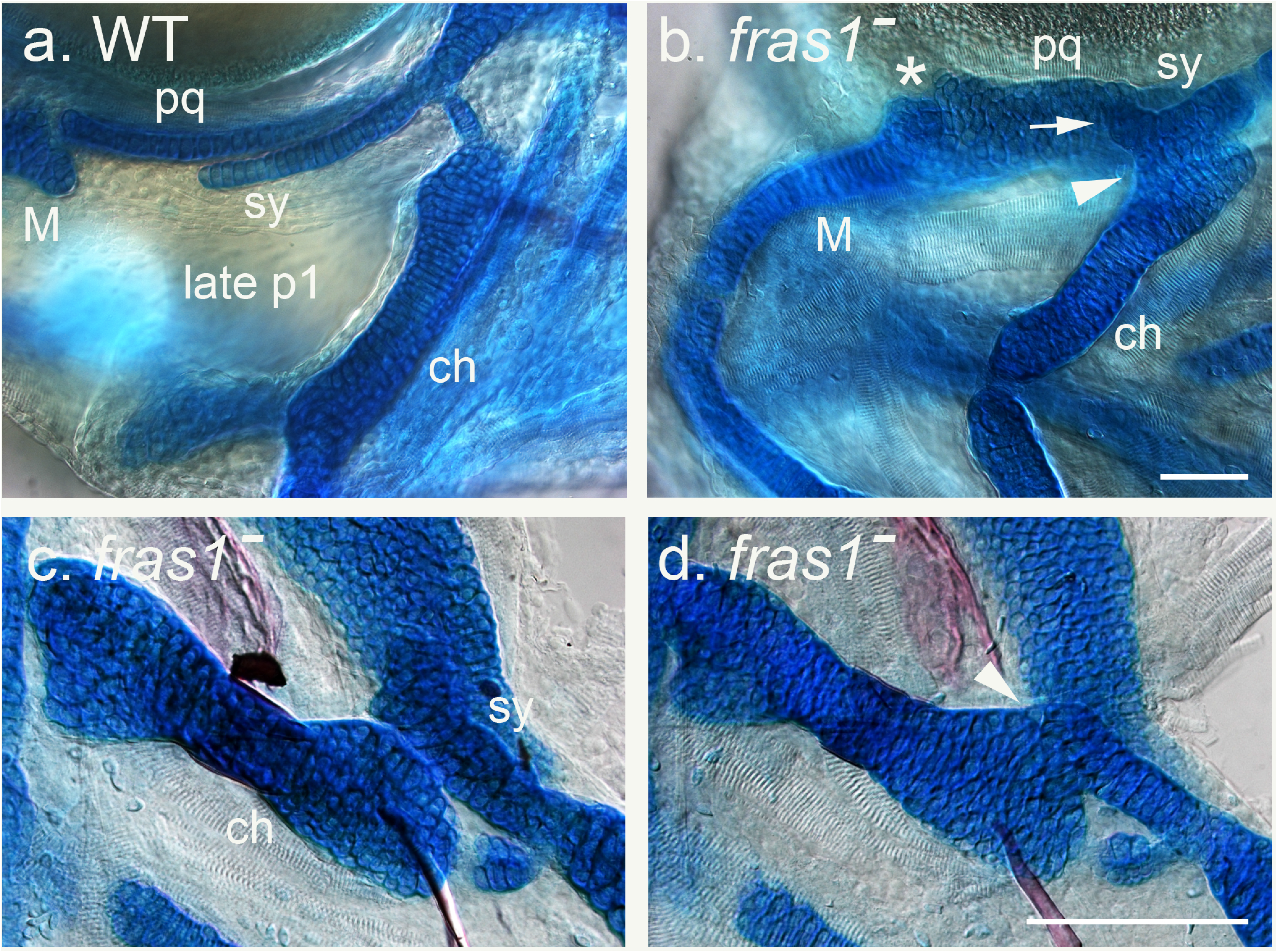
Details of larval skeletal anatomy; Alcian Blue-Alizarin Red stained preparations, photographed using Nomarski interference contrast illumination to enhance contrast and reveal unstained tissues. Similar orientations to Figure 1. (a) Wild type, showing the cartilages surrounding the late-developing region of the first pharyngeal pouch (late p1). Cartilages are labeled as in Figure 1. (b-d) *fras1* mutants. (b) Two mutant fusions are illustrated, the M-pq (asterisk) and sy-ch (arrowhead). The sy has a short unfused extended region about three-four cells long (arrow), distal to the fusion. A band of muscle (striations visible) is present where late p1 would be located in the WT. (c, d) Higher magnification views of sy-ch configurations. (c) the sy and ch come into close apposition, but we do not score this condition as a full fusion; note the thin blue line separating the two elements. (d) Here, in contrast to c, the sy-ch full fusion is evident from the way that the cartilage cells from the two elements are interdigitated. In contrast to b, there is no extended region of the sy in this case.

### 1.3 Role of the endoderm

Epithelia specifically express the *fras1* gene, whereas the cartilages form from neural crest-derived mesenchyme that does not express *fras1* (Talbot *et al*., 2012). Hence, we can infer that the skeletal phenotypes due to loss function of *fras1* are indirect (nonautonomous) at the cellular level. This inference has experimental support: Talbot *et al*. (2012) transplanted cells between wild-type (WT) and *fras1* mutant embryos to make genetic mosaics, revealing that in genetic mosaics only the pharyngeal endoderm need to be mutant in genotype for the mesenchymally derived cartilages to express the mutant fusion phenotypes. Reciprocally, transplantation of WT endoderm can rescue the cartilage phenotypes in otherwise mutant individuals. In agreement with the expression studies, these genetic mosaic experiments show that the *fras1* skeletal phenotype is cell-nonautonomous in the cartilage.

What is the nature of epithelial (endodermal) - mesenchymal interactions that provide for proper cartilage morphogenesis in WT embryos? Talbot made two key discoveries that provide insight into this question. First, he discovered that a portion of the first endodermal pouch, he named “late p1” (Figure 2A) does not form in mutants (Talbot *et al*., 2012, and see Figure 2). Second, late p1 normally forms by outpocketing from the deep medial endoderm toward the outer ectoderm at just the stages that early cartilage morphogenesis is getting underway in the immediate vicinity of the pouch. Furthermore, timed heat shocks with a conditional *fras1* allele revealed just when the WT gene function must be present in the endoderm for p1 to form, and for WT cartilage morphogenesis: The *fras1* critical period coincides closely with late p1 outpocketing (Talbot *et al*., 2016). The strong suggestion from this work is that late p1 provides a key component of the embryonic environment specifically during the early period of skeletal morphogenesis. It is possible (but as yet unproven) that cartilage precursors use the pouch endoderm, and in particular, its associated Fraser protein complex, as a cellular/molecular pathway for morphogenetic outgrowth of the cartilages. We note an important caveat is that phenotypic variation of the cartilage fusions does not depend on late p1 in a one-to-one manner because failure of late p1 outpocketing is fully penetrant in *fras1* mutants, but cartilage mutant phenotypes are not. Furthermore, studies of embryos at different ages showed that some of the mutant fusions only form well after the time when pouch outpocketing would normally occur (Talbot *et al*., 2012). Full understanding will require more work.

### 2.4 Off-target action of *Gal4VP16*

In the course of time-lapse experiments using the widely known Gal4-UAS system to drive GFP fluorescence in neural crest derived cranial mesenchyme, we discovered, and report below, an unusual “off-target” action of the transgene *fli1a-F-hsp70l:Gal4VP16*. We see enhanced penetrance of the mutant skeletal phenotypes in *fras1* mutants. The action of the transgene is off target in that we show the enhancement is present in the absence of the normal Gal4 target, *UAS*. Finding of an off-target effect of a *Gal4VP16* transgene is not original. The *Gal4VP16* was engineered in the laboratory of M. Ptashne and its use encouraged because it is very potent transcriptional activator (Sadowski *et al*., 1988). The same group quickly discovered that transcription of off-target genes (i.e., genes other than *UAS*) could be inhibited, rather than promoted, by high activity of *Gal4VP16*. The effect, they termed “squelching” was proposed to be due to sequestration (titration) of a transcription factor due to the unusually high activity of the transgene (Gill & Ptashne, 1988). Furthermore, Koster & Fraser (2001) described toxic effects of *Gal4VP16* (which could be due to squelching) including generalized cell death and abnormal development in zebrafish embryos. Toxicity of *Gal4VP16* has been reported by several groups (Koster & Fraser, 2001; Scott *et al*., 2007; Asakawa *et al*., 2008), and led workers in the Kawakami laboratory to engineer a new version, *Gal4FF*, that they showed was substantially less toxic than *Gal4VP16*, but still showed considerable activation of *UAS* (Asakawa & Kawakami, 2009). Toxicity led other workers to abandon the Gal4-UAS system altogether, for example in favor of a new transgenic “Q” system to express genes of interest (Ghosh & Halpern, 2016). Squelching and/or toxicity could be responsible for the enhancement we describe, as we consider in the Discussion.

## 2 MATERIALS AND METHODS

### 2.1 JFB Ethics statement

No fishes were collected as part of faunal surveys. Zebrafish were killed during the experiments. No surgical procedures were performed. No experimental conditions severely distressed any fishes. No procedures caused lasting harm to sentient fishes. No procedures involved sentient animals subjected to chemical agents.

### 2.2 Fish husbandry and strains

All of the studies were done with zebrafish of the strain AB genetic background, developed and maintained at the University of Oregon as a closed strain for over 40 years. The fish were kept at 28.5°C, raised, maintained and crossed as described, (Westerfield, 2007). All procedures with live fish followed approved University of Oregon IACUC guidelines. The *fras1^te262^* mutant, likely a null (Carney *et al*., 2010, Talbot *et al*., 2012) was originally isolated on the TU background. We crossed it into AB, and used this allele for the experiments reported here. The *fras1* homozygotes are usually lethal and we carried the line with heterozygotes, outcrossed each generation to AB. Heterozygotes were identified by PCR genotyping, following Carney *et al*. (2010) and Talbot *et al*. (2012), with primers GGAAGATTTTCTTTATTTAGCAGTCTCT and TTGGAACTAGGTCCTCTTTGGTGTGCTATAAAATA, and digestion with SspI. We identified homozygous mutant embryos and larvae from the fully penetrant fin blistering phenotype (Carney *et al*., 2010) that we score at about 48 hours postfertilization or later. The *fli1a-F-hsp70l:Gal4VP16*^*el360*^ transgene, which carries the *my17:EGFP* transgenesis marker driving fluorescence in the heart, and the *UAS:GFP* transgene were gifts, as stable integrations, from the G. Crump lab (Das & Crump, 2012). The *fli1a-F-hsp70I* promotor drives high expression of *UAS:GFP* in the pharyngeal arch neural crest-derived ectomesenchyme (specificity due to the *fli1a*-F), and also low levels of expression in several other tissues of the body, the latter probably due to the minimal heat-shock promotor (hsp70I). The *fli1a-F-hsp70l:Gal4VP16* transgene and the *UAS:GFP* transgene are unlinked to one another and unlinked to *fras1*. The mutant and transgenic lines were maintained by outcrossing to AB, although incrosses were used in particular experiments, as described in the RESULTS section. Live embryos were screened with a fluorescence dissecting microscope for GFP fluorescence in the heart, indicating presence of the *fli1a-F-hsp70l:Gal4VP16* transgene, and for whole body GFP fluorescence, especially in the pharyngeal arches, indicative of the presence of both the *fli1a-F-hsp70l:Gal4VP16* and *UAS:GFP* together. The larval skeletons, at 7 days postfertilization, after euthanization and brief fixation, were stained with Alcian Blue and Alizarin Red, as described (Walker & Kimmel, 2007). Acridine Orange staining of live whole-embryos, was carried out for 30 minutes, using 1 µg/ml or 5 µg/ml in Embryo Medium at 48 hours postfertilization, essentially as described (Tucker & Lardelli, 2007). Fluorescence was examined with GFP filter sets

We also obtained the *fli1a-F-hsp70l:Gal4VP16*:pA_CG2 plasmid from the G. Crump lab, that they had used to make our transgenic line ‘A’. We micro-injected 0.5 nl of injection mix containing this plasmid (30ng/µl) with the Tol2 RNA transposase (20ng/µl) into 1-cell zygotes from a cross of *fras1* heterozygotes, and not bearing the integrated *fli1a-hsp70I:Gal4VP16* transgene to study possible enhancement due to mosaic expression of the transgene, and to make new, independent permanent lines. In one of two experiments, one of the parents of the recipient fish was also heterozygous for the *UAS:GFP* transgene, in order to be able to assess whether the injected Gal4 was able to activate *UAS:GFP* in the pharyngeal arch mesenchyme of the mosaic recipients.

### 2.3 Transgene mapping using genomic NGS libraries

Genomic DNA was extracted from muscle tissue of one individual carrier fish from each of the transgenic lines using the DNAeasy kit (Qiagen). One microgram of genomic DNA was sheared to an average length of 475 base pairs (bp) using a Covaris M220 Sonicator. Sonicated DNA was used to construct genomic libraries using the Kapa Hyper Prep Kit following manufacturer’s recommendations, and using dual-indexed Truseq Illumina sequencing adapters provided by the University of Oregon Genomics and Cell Characterization Core Facility. Genomic libraries were double-size-selected with AMPure XP beads to an average size of 500 bp, then amplified with four cycles of PCR. Amplified libraries were cleaned up with AMPure XP beads, normalized to 3 ng/ul, and multiplexed. Next generation sequencing was performed on an Illumina Hi Seq 4000 using paired-end 150 bp reads. Adapters were removed with Cutadapt and reads were quality-trimmed with Trimmomatic and aligned using GSNAP to the zebrafish genome assembly GRCz11 (Bolger et. al., 2014; Howe et al., 2013; Wu et al., 2016).

### 2.4 Phenotypic scoring of the pharyngeal skeletal phenotypes

Alcian Blue and Alizarin Red-stained larvae, in 50% glycerol-0.1%KOH, were distributed individually into the wells of 96 well plates, the wells coded such that scoring could be done ‘blind’ as to *fras1* genotype and whether fluorescence due to the *fli1a-F-hsp70l:Gal4VP16* transgene and/or *UAS:GFP* transgene were present in each individual specimen. The left sides and right sides of the fish in these wells were blind-scored on different days, with care not to refer to the side (or data) not being scored. Additionally, several of the experiments were blind-scored independently by two observers, allowing us to estimate ‘technical error’ due to scoring problems.

For the five traits described in the Results section, we uniformly use categorical variables, and binary coding; a score of 1 is the mutant condition, and a score of 0 is the wild-type or near wild-type condition. To score an individual larval phenotype, a larva was removed from its well and placed in a 30 mm culture dish containing glycerol-KOH, so that positioning was easy to accomplish to allow the best views of the different traits. Scoring was done with the highest magnification achievable with a standard Zeiss Stemi 2000 dissecting microscope, equipped with a 2x extension lens. Three cartilage fusions were scored, fusions between 1) the symplectic (sy) and ceratohyal (ch) cartilages, 2) the symplectic and the palatoquadrate (pq) cartilages, and 3) the pq and Meckel’s cartilage (M). For each trait, we recorded whether a fusion was present – i.e., we could see cartilage cells from one of the partners well integrated with cells from the other (as in Figs. 1 and 2), versus a close apposition of the two elements with no fusion (e.g., as in Figure 2C), versus a wild-type looking condition. For the data reported here the latter two categories were combined, so that the scores, for each fusion, could be reduced to 0 (no fusion) or 1 (fusion). In some experiments, for the sy-ch fusion we also recorded whether distal sy cartilage cells were present that extended past the site of the fusion, and if so, how many cells were present in the extended region. For the plots reported here, these data were reduced to 0 (sy extension present – the milder phenotype) or 1 (not present). Part way through the course of the study we observed severe malformation of the opercle bone (op), and in the later experiments we recorded whether the op had the fan shape characteristic of the normal young larval stage, a reduced fan (as in Figure 1C), for the mutant condition, or a ‘stick’ condition of various lengths, where no broadening out of the distal end of the bone was present. For the plots, these data were reduced to 0 (fan shape) and 1 (the more severe stick shape). As explained above, these data reductions resulted in two categories (0 or 1, indicating WT-mild or severe respectively) for each of the five traits, (sy-ch, sy-pq, M-pq fusions, sy extension, op shape).

For analysis, the data were entered into JMP (SAS Institute Inc., v. 13) as categorical (non-numeric) variables. Mosaic plots and contingency tables were computed. Determination of significant differences between groups used the Chi-square distribution, and the Pearson estimation of probabilities, available in JMP. Differences were considered significant at alpha=0.05. Developmental instability was estimated from unilateral expression-frequency data as described (Hallgrimsson *et al*., 2005).

## 3 RESULTS

### 3.1 The *fli1a-F-hsp70l:Gal4VP16* transgene enhances penetrance of the fusion phenotype

In our studies, using the Gal4-UAS system to express GFP in craniofacial cartilages, we noted that the cartilage fusions described above for *fras1* mutants showed enhanced penetrance among the transgene-bearing *fras1* mutants. Moreover, the penetrance of a previously undescribed fusion, between the sy-pq, was also enhanced in the transgenic mutant larvae. These three fusions, as well as two other skeletal traits we examine below, are essentially unique to the homozygous *fras1* mutants: Phenotypically wild-type siblings, two-thirds of which are expected to be *fras1* heterozygotes, show only background levels of the traits, whether or not they were transgenic, with penetrances of 0-3.6% (Table 1). Below, we ignore the wild types, and examine the traits in transgenic and control (non-transgenic) *fras1* mutants. Technical error, estimated by determining the difference in blind scoring between two independent investigators, was of the order of 5-10% for three of the cartilage traits scored, and considerably less for the sy-pq fusion (Table 2).

To investigate heritability of the enhancement in mutants, we grew up and intercrossed phenotypically wild-type siblings of the *fras1* mutants (two-thirds of which are expected to be *fras1* heterozygotes) that we knew were positive for both the *fli1a-F-hsp70l:Gal4VP16* and the *UAS:GFP* transgenes because their hearts and pharyngeal arches expressed GFP fluorescence. Results from the F1 progeny of this cross verified heritability, in that the transgene-possessing mutants showed significantly higher penetrance for all three cartilage fusions than mutant siblings without either transgene (Figure 3). Furthermore, the penetrance of the mutants with the *fli1a-F-hsp70l:Gal4VP16* by itself, and mutants with both *fli1a-Fhsp70l:Gal4VP16* and *UAS:GFP* showed no significant differences (Figure 3). This analysis importantly showed that the *UAS:GFP* transgene apparently played little or no role in the enhancement, and that the transgene of interest was *fli1a-F-hsp70l:Gal4VP16*. The penetrance plots for the left and right datasets look very similar to one another, lending confidence that the scoring was accurate. (Completely identical plots are not expected, differences between the sides arise due to developmental instability, discussed below). Because, as we show in this experiment, the *UAS:GFP* is apparently irrelevant with respect to the enhanced *fras1* mutant phenotypic severity, we made a new (F2) generation, selecting parents from the F1s that bore the *fli1a-F-hsp70l:Gal4VP16* transgene but not the *UAS:GFP* transgene (i.e., we crossed the *UAS:GFP* out of this line, that we call “line A”, to simplify comparison with other lines described 7 below). This selection was possible because progeny expressing heart fluorescence only will have not inherited *UAS:GFP*.

**FIGURE 3.**
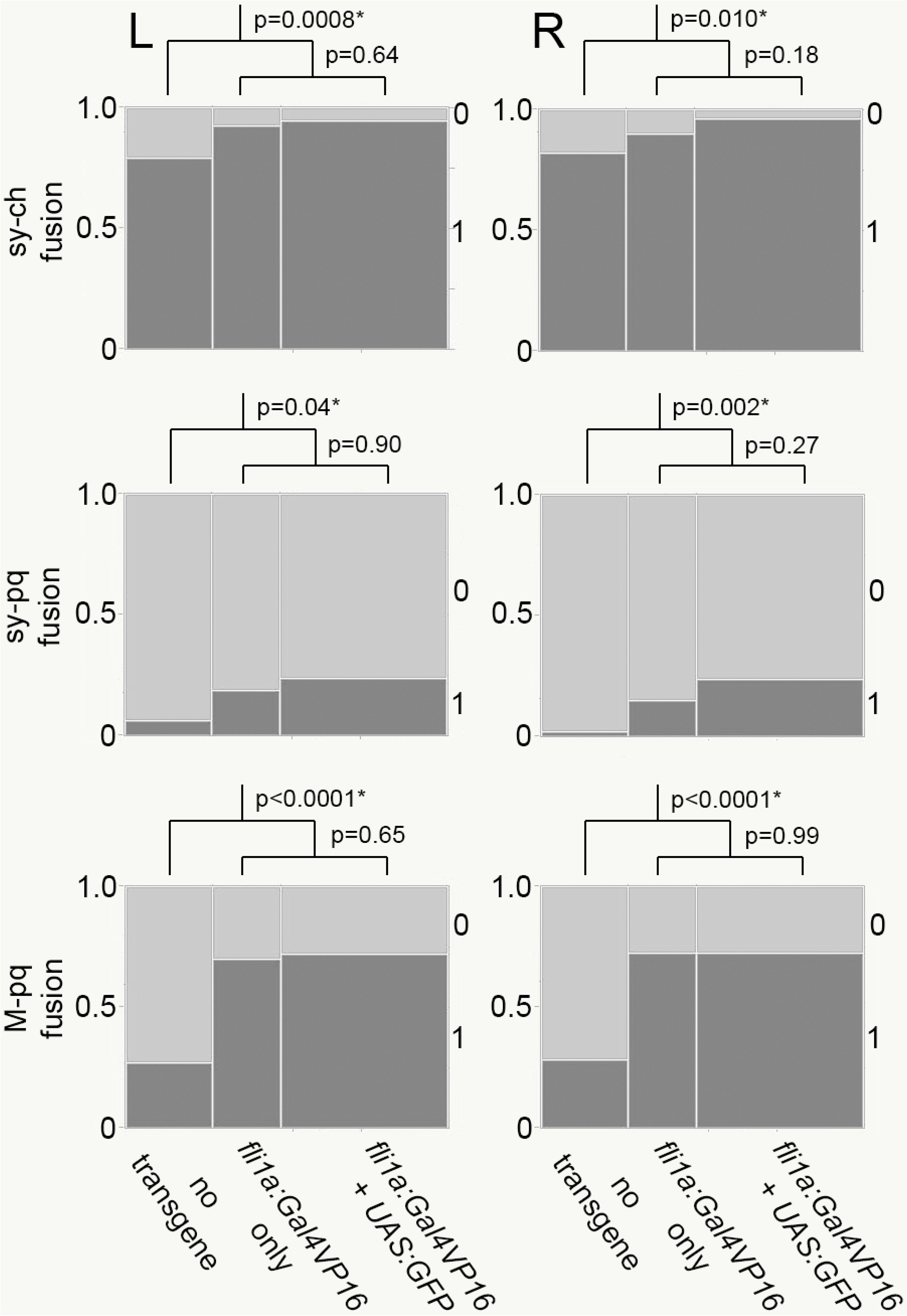
Heritable increase in penetrance in three cartilage fusion phenotypes due to presence of the *fli1a-F-hsp70l:Gal4VP16* transgene. Mosaic plot comparing *fras1* mutant siblings, with and without transgenes. Normalized penetrance of the fusion is indicated by the lower, more darkly shaded part of a bar (y axis, range from 0 to 1). The numbers along the right side of each plot indicate the presence (1) or absence (0) of the mutant phenotype. The normalized fraction of each transgene class is given by the width of the bars. Scores from the left and right side of the animals are shown as separate plots (L, R). The p-values comparing the two transgene classes to each other (lower for each plot) showed no significant difference for all of the plots at apha=0.05. The p-values comparing the mutants with no transgenes to the combination of both transgene classes together (upper for each) showed significant differences for all of the plots. A set of 188 mutants were scored, obtained from incrosses of parents each heterozygous for fras1, and one parent heterozygous for a single copy of the *fli1a-F-hsp70l:Gal4VP16* transgene, and heterozygous for the *UAS:GFP* transgene. Progeny were scored as *fras1* homozygous mutants by the blistered fins phenotype, and scored by fluorescence for presence of neither transgene (no fluorescence) or either the *fli1a-F-hsp70l:Gal4VP16* transgene alone (fluorescent hearts only), or the combination of the *fli1a-F-hsp70l:Gal4VP16* and *UAS*:*GFP* transgene (fluorescent hearts, bright pharyngeal arches and lower general body fluorescence).

### 3.2 Enhancement by *fli1a-F-hsp70l:Gal4VP16* appears dose-dependent

In the experiments described above we used within-strain crosses, *in*crosses of *fras1*^−^ heterozygotes bearing single copies of the transgenes, to obtain progeny that could be examined for fusion penetrance. With incrosses, the *fli1a-F-hsp70l:Gal4VP16* transgene could be inherited from neither, either, or both parents, which would result in copy number being 0, 1, or 2 in individual progeny. We compared severities of homozygous *fras1* mutant larvae obtained from such incrosses (i.e., both parents have the genotype *fras1^+/−^ fli1a-F-hsp70l:Gal4VP16*) with progeny obtained from *out*crossing a fish of this genotype to a *fras1* heterozygote that did not contain the transgene (i.e., *fras1*^+/−^ *fli1a-F-hsp70l:Gal4VP16* crossed to *fras1*^+/−^). Progeny from this outcross will have only 0 or 1 transgene copies, not 2.

The incross and outcross progeny both showed enhanced penetrance of *fras1* traits due the presence of the *fli1a-F-hsp70l:Gal4VP16* transgene, as compared with their mutant siblings that did not possess the transgene (Figure 4). The data for the outcross establish that the transgene acts as a dominant in terms of its enhancer function: This is because all of the transgene-bearing outcross larvae will possess only a single transgene copy.

**FIGURE 4.**
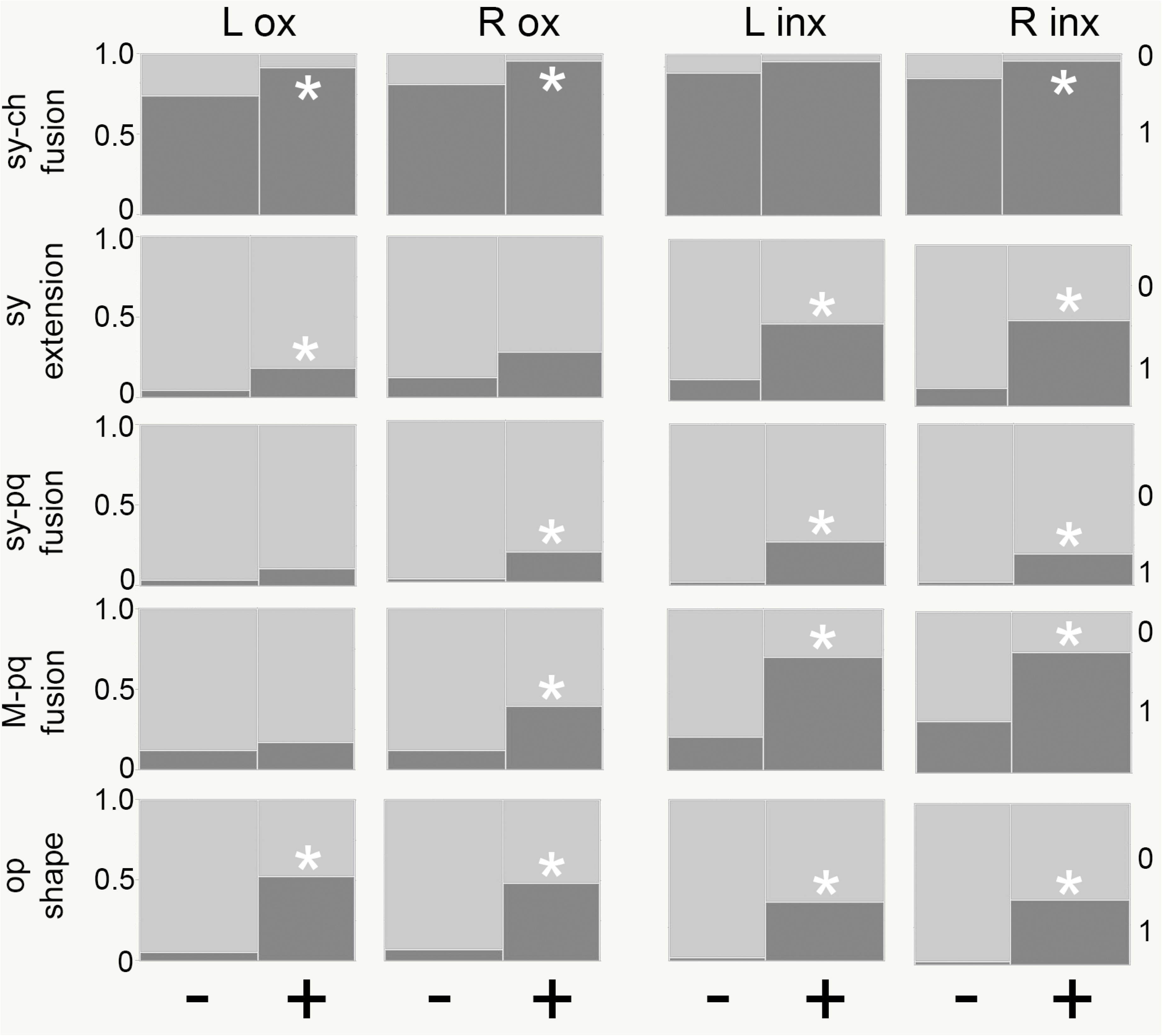
For progeny of both incrosses (inx) and outcrosses (ox) most traits show significantly higher penetrance when the *fli1a-F-hsp70l:Gal4VP16* is present (+) than when no transgene is present (−).Mosaic plots as in Figure 3. Significance at alpha=0.05 is indicated by an asterisk. Increased penetrance of the transgene-bearing outcross progeny, as compared with progeny without the transgene, indicate that a single transgene copy is sufficient, i.e., the transgene behaves as a dominant trait. n=120 individuals for the inx progeny and 106 for the ox progeny. Incross data are from crosses as explained in the legend to Figure 3 (but here *UAS:GFP* is not present). Outcross data are from crosses in which one parent only was heterozygous for presence of the *fli1a*-F-hsp70l:Gal4VP16; both parents were heterozygous for *fras1*^−^.

Comparisons, of progeny from incrosses and outcrosses that all possessed the transgene (one or two copies for the incross) are shown in Figure 5. The sy-ch fusion scores were insignificantly different for the two groups (i.e., incross versus outcross), seemingly because in this experiment the penetrance scores were very close to 100%, which would make distinguishing the two groups difficult. To further explore the severity of the sy-ch fusion we included another, related, trait: namely whether an unfused distal tip of the sy extended beyond fusion with the ch (scored as ‘0’; see Materials and Methods), versus no extension being present (the more severe phenotype, scored as ‘1’). Figure 5 shows a significant difference in penetrance of the sy extension between the incross and outcross on both the left and right sides: the incross mutants have significantly higher penetrance than outcross progeny. Furthermore, matching this finding, *fras1*^−/−^ transgene-bearing cross progeny showed substantially higher penetrance for the M-pq fusion phenotype on both sides of the body than *fras1*^−/−^ transgene-bearing outcross progeny (Figure 5). We interpret these higher penetrances in the incross progeny for the sy extension and the M-pq fusion to mean that two copies of the transgene (expected to be present in a third of the incross progeny, and not at all in the outcross progeny) impart higher severity than a single copy. We conclude that, for some phenotypes, the transgene behaves genetically like a partial dominant.

**FIGURE 5.**
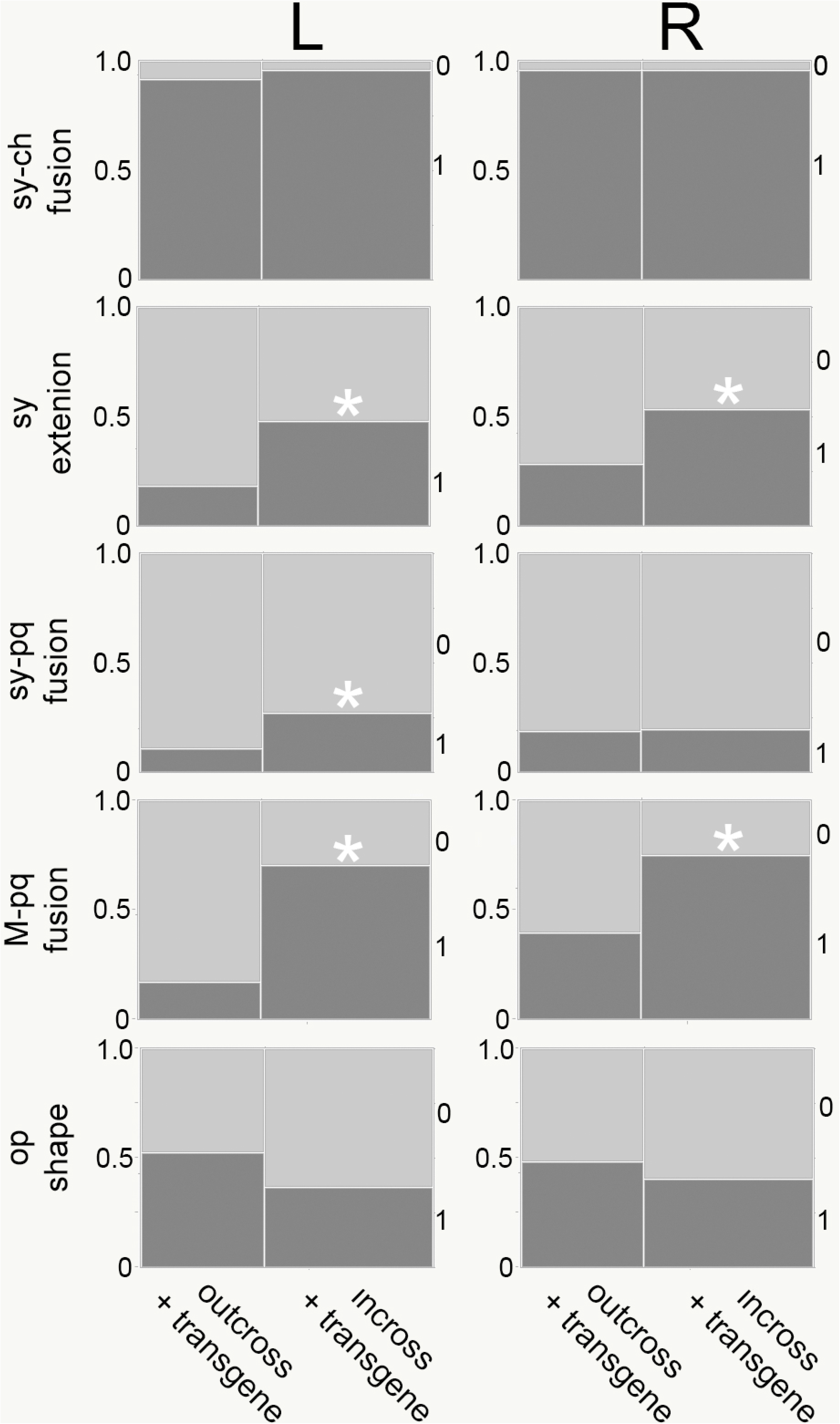
For some *fras1* mutant traits, particularly the sy extension and the M-pq fusion, *fli1a-F-hsp70l:Gal4VP16* transgene-bearing incross progeny show significantly higher penetrance than transgene-bearing outcross progeny. Mosaic plots as in Figure 3, here comparing incross and outcross groups. We use only the *fli1a-F-hsp70l:Gal4VP16* transgene classes for this analysis. Significant differences are indicated by asterisks. n=115 individuals. Incross data from crosses as explained in the legend to Figure 3 (but here *UAS:GFP* was not present). Outcross data from crosses in which one parent only was heterozygous for the *fli1a-F-hsp70l:Gal4VP16* transgene; both parents were heterozygous for *fras1*^−^.

In the course of these experiments we also noted another previously undocumented *fras1* mutant phenotype – a change in shape of the opercle bone, from the usual ‘fan’ shape (scored as ‘0’) characteristic of the early larva to a ‘stick’ shape, more severe (scored as ‘1’). As shown in Figure 4, the stick shape is present but very rare in non-transgene possessing *fras1* mutants, and penetrance is substantially (and significantly) increased when the transgene is present. However, the penetrance is about the same between the incross and outcross transgene-bearing groups (Figure 5).

The incross-outcross experiments also provide us with an opportunity to test the hypothesis that toxic effects of the *fli1a-F-hsp70l:Gal4VP16* explain the higher phenotypic penetrances of transgenic mutants. As noted in the Introduction, toxic effects have been associated with high expression of Gal4VP16 (Koster & Fraser, 2001; Scott *et al*., 2007; Asakawa *et al*., 2008). Hence the increased penetrance of *fras1* mutant phenotypes in the presence of the transgene could be due to dead or dying cells, in spite of our not detecting any over-all deformities or evidence of cell death. To test this proposition, we stained incross mutant larvae with Acridine Orange, with and without the transgene, to detect apoptosis, a possible consequence of toxicity. The highly reduced and blistered fins in the *fras1* mutants fluoresced brightly with this stain, whereas the fins of wild-type larvae showed only little fluorescence, as did the rest of the body. Other than the fins, Acridine Orange staining revealed no difference between wild types and mutants, and whether or not the *fli1a-F-hsp70l:Gal4VP16* transgene was present. In particular, we could not pick out a subset of the transgene-bearing mutants within the incross progeny that had more Acridine Orange labeling, those with two copies expected to be present in a 1:3 ratio, as compared with those with a single copy of the transgene. Thus, Acridine Orange staining yielded no evidence for the increased penetrance of *fras1* mutant phenotypes being due to toxicity.

These data provide clues as to the nature of the genetic interactions between the *fli1a-F-hsp70l:Gal4VP16* transgene and *fras1*. The data show that for all the mutant traits examined, a single chromosomal copy of the transgene is sufficient for enhanced penetrance, and further suggest that for two of the traits, two copies further increase penetrance over the single copy. Dominance implies that there is something special about inheritance, perhaps a gain of function, as compared with a recessive (loss of function) phenotype. Partial dominance further implies that if there is such a ‘new’ function dependent at least indirectly on the transgene, it can be expressed at greater or lesser extents: transgene dosage matters.

### 3.3 The site of integration of the transgene is not the critical determinant of enhanced severity

We can consider that the increase in severity of aspects of the *fras1* mutant phenotype is an ‘off-target’ action of the *fli1a-F-hsp70l:Gal4VP16* transgene, in the sense that the ‘on-target’ function is the activation of *UAS* of a second transgene, which is not present in our A line mutants, since *UAS:GFP* was crossed out of the A line. A straight-forward interpretation of the dominance results just described is that sequences within the transgene itself are causal in enhancing the *fras1* mutant phenotype and likely in a dosage-dependent manner. The observation, however, does not rule out an alternative model, that a change in a host genomic sequence located adjacent to or near the transgene integration site is critical. Insertional mutagenesis is a well-known phenomenon depending on local host sequences, and can explain many phenotypic changes due to transgenes. Clearly the site of integration is critical for insertional mutagenesis, but integration site is expected to be less important if the enhancement were primarily due to transgene sequences themselves.

To test the hypothesis that the site of integration is critical for enhanced *fras1* phenotypic severity we injected, into *fras1* mutants, Tol2 transposase and the *fli1a-F-hsp70l:Gal4VP16*-containing plasmid used to make the *fli1a-F-hsp70l:Gal4VP16 insertion* for our original line – line A, asking if increased *fras1* phenotypic penetrance occurred in transient mosaics. There was no enhancement of penetrance of the three cartilage fusion traits in two rounds of injections (Figure 6). Taken at face value, our finding no enhancement in the mosaics could mean that an integration site is critical, but another possibility is that even though mosaic GFP fluorescence was easily detectable in recipients of the plasmid, showing presence of the transgene, the level of activity of a putative causal transgene sequence, or the number of cells possessing the transgene, in the injected mutants was insufficient for enhancements. To address these possibilities, we selected for new stable independent lines by rearing injected, phenotypically wild-type siblings of the injected *fras1* homozygous mutants. Since the plasmid has no sequences directing integration to a particular genomic location, we assumed that the integrations would occur essentially at random with respect to host sequences – at the very least, new insertions are not expected to be at or near the insertion site of the original line A. We obtained five new stable transgenic *fras1* mutant lines in this experiment, all independently derived, possessing the *fli1a-F-hsp70l:Gal4VP16* transgene. We first mapped our original insertion line A toward the right end of *Danio rerio* chromosome 20 (*Dre*20:41,998,502-42,203,629). As expected, all of the new lines appeared to map to different chromosomes: line B (Dre25:34,323,250), line C (Dre4:41,046,100), line D (Dre21:13,196,000), line E (Dre2:10,950,400), line G (Dre22:18,305,000). We conclude that all of our new lines went into locations different from the original line and different from one another. Figure 7 compares the penetrance scores of *fras1* mutant traits between the transgene containing (+) and non-containing (−) *fras1* mutant progeny derived from line A, and each of the new lines (B-G). We detected no significant enhancement due to the transgene in line D, however the other new lines (B, C, E, G) showed significantly enhanced severity for at least one trait when the transgene was present. This majority result shows that sequences within the transgene rather than the site of integration are the critical determinants of the enhancement.

**FIGURE 6.**
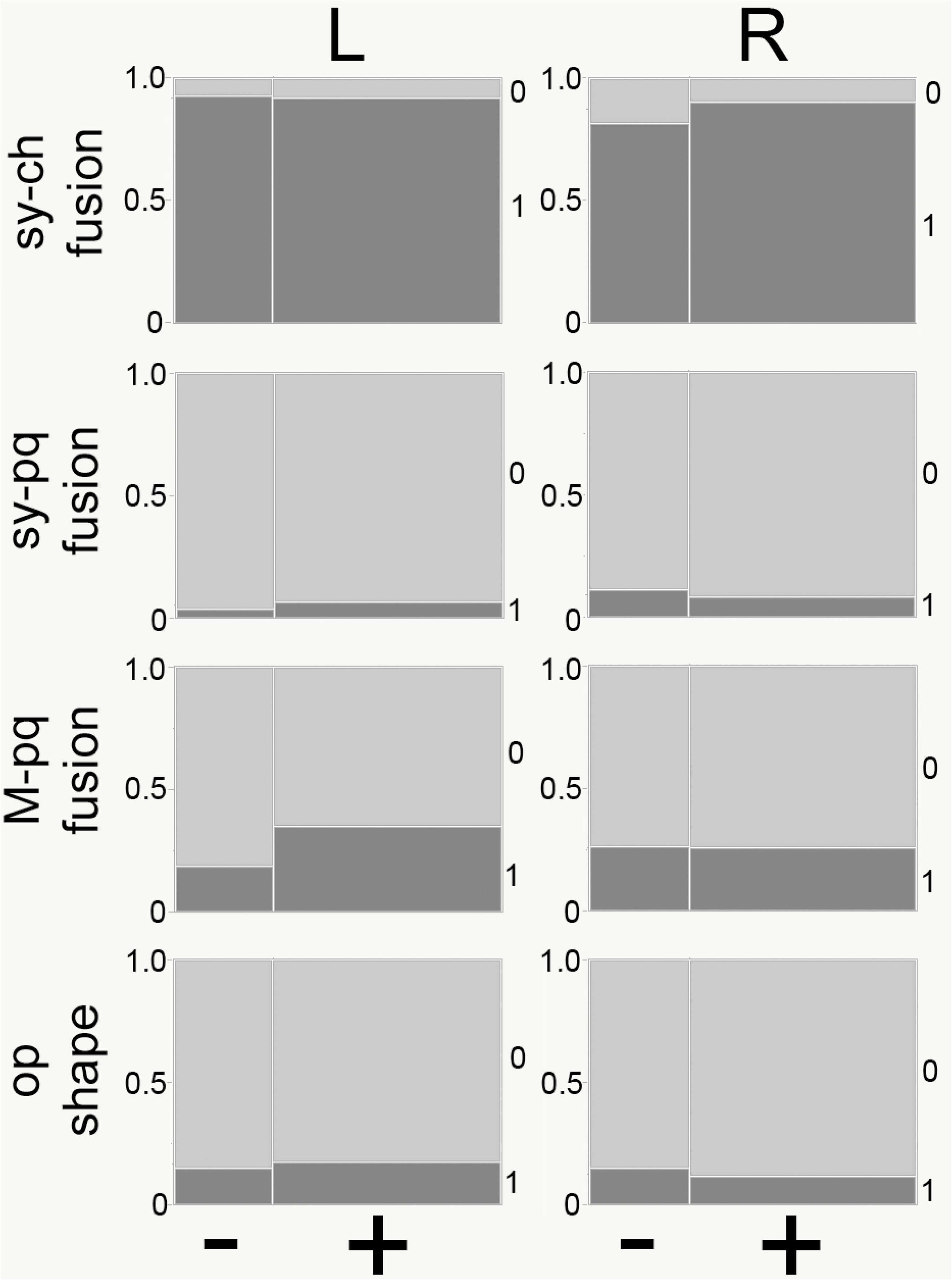
Micro-injection of the *fli1a-F-hsp70l:Gal4VP16*-plasmid results in no significant enhancement *of fras1* mutant phenotypes in the recipients. Mosaic plots as in previous figures. In the experiment shown, the plasmid was injected into eggs derived from fish heterozygous for *fras1*^−^, and heterozygous for *UAS:GFP*. The resulting embryos were scored for *fras1*^−^ homozygosity, for fluorescence in the heart and pharyngeal arches, and, after skeletal staining, for *fras1* mutant skeletal phenotypes. The comparisons shown are between classes with medium-to-high arch fluorescence (+), and fluorescence (−); individuals with only dim fluorescence were omitted from the analysis). None of the comparisons showed a significant effect of the transgene in these ‘transient’ mosaic analyses.

**FIGURE 7.**
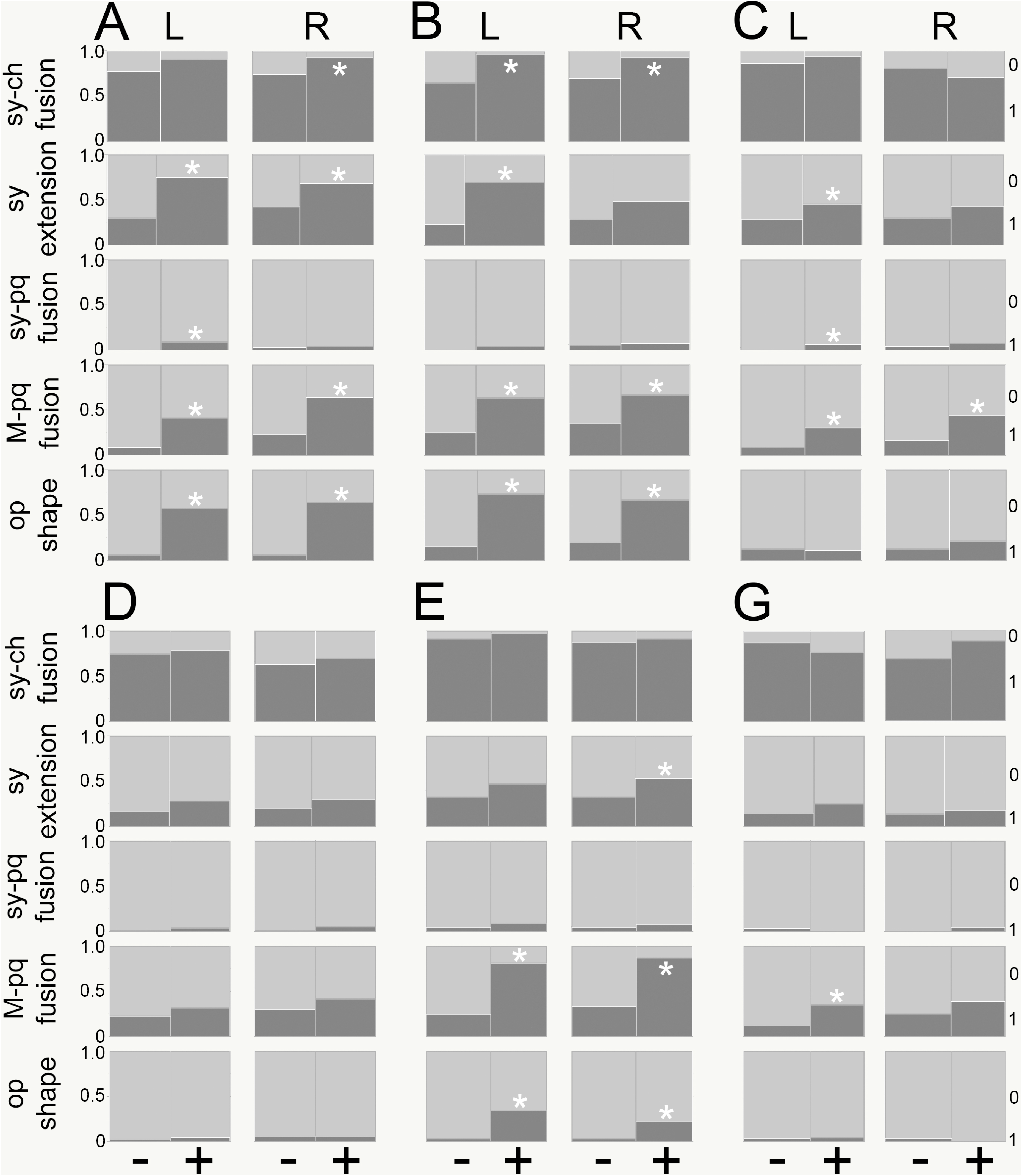
Four out of five new, independently derived *fli1a-F-hsp70l:Gal4VP16* transgenic lines (lines B-G) enhance penetrance of at least some *fras1* mutant phenotypes. F1 progeny from the plasmid injected fish. Line A is our original line. n=712 individuals. Mosaic plots as in previous figures, comparing penetrance in the presence and absence of the transgene. Asterisks indicate differences significant at alpha=0.05. Enhancement of penetrance varies remarkably among the new lines – B is approximately as active as line A, and line D showed no significant enhancement (see also observations described in the text). For the B-D lines no *UAS:GFP* is present. Gal4 function was assessed In line D, and found to be present, by raising F1s and crossing the transgenic adults to a heterozygous *UAS:GFP* stock. For lines E and G, the recipients of the plasmid possessed the *UAS:GFP* transgene, such that we could show directly that Gal4 was functional in the F1s.

### 3.4 Prominent phenotypic diversity among independent *fli1a-F-hsp70l:Gal4VP16* transgenic lines

The data show considerable variation in phenotypic penetrance among the lines just described. As mentioned earlier in the Results section, the considerable similarity between corresponding scores on the left and ride sides of the same set of mutants strongly suggests that the variation we encountered in this experiment is not due to inaccurate scoring, done blindly and the left and right sides scored on different days. Among the data shown in Figure 7, line E has the largest penetrance score for the M-pq fusion we observed in the presence of the transgene, (81% for the left side) in this experiment, and the largest difference between scores when transgene is present or absent, (57%). In contrast, line C, showed the smallest score for this trait, the corresponding values are 30% and 22%, with and without the transgene, respectively. Penetrance varies among the lines across the matrix. Line A shows significant differences between transgenic and nontransgenic individuals for eight out of the ten traits (considering the left and right sides separately). Line B is close to line A, with seven of the ten comparisons showing significance. Line E has five significant differences, line C four, and line G a single one. Line D had no significant differences at all. Which traits appeared significantly different among the lines appeared to be uncorrelated. The most robust of the four phenotypes of those examined, i.e., being resistant to perturbation by the transgene, are the sy-pq fusion, showing significant differences in two of the twelve comparisons, and the sy-ch fusion, showing significant differences in three (all of these restricted to lines A and B). The least robust is the M-pq fusion, with significant differences in nine. The sy extension and op shape phenotypes are intermediate, with five and six significant differences respectively. Interestingly, for the sy extension we see several cases where one side shows significance and the other doesn’t (lines B, C, E), but for the op phenotype, only left right pairs of significant differences are observed (lines A, B, E). Phenotypic variation is surprisingly high among these transgenic lines, considering that they were all derived from the same injections with the same transgene.

### 3.5 Possession of the *fli1a-F-hsp70l:Gal4VP16* does not appear to increase developmental instability

Increased phenotypic variation can arise from increased developmental instability (DI), stemming from loss of buffering of stochastic features of development (Polak, 2003; Dongen, 2006; Graham *et al*., 2010). DI of a trait can be estimated from the categorical (non-numeric) variables we use in this study, by examining unilateral expression of the trait, taken to be a feature of DI. Hallgrimsson *et al*. (2005) showed that trait frequency must be taken into account to draw valid conclusions about DI, and, following these authors we calculate the proportion of unilateral expression, U, as function of trait frequency, F. The slope of this relationship (U vs. F) is related to DI, with a greater slope indicating more DI (Hallgrimsson *et al*., 2005). Here we use penetrances of the traits, taken from the large dataset shown in Figure 7. We compare all the traits we examined in transgene-possessing larvae with no-transgene possessing larvae irrespective of their genetic lines. Figure 8 shows that slopes for transgene-possessing larvae and no-transgene possessing larvae are essentially identical, thus providing no evidence that possession of the transgene has increased developmental instability.

**FIGURE 8.**
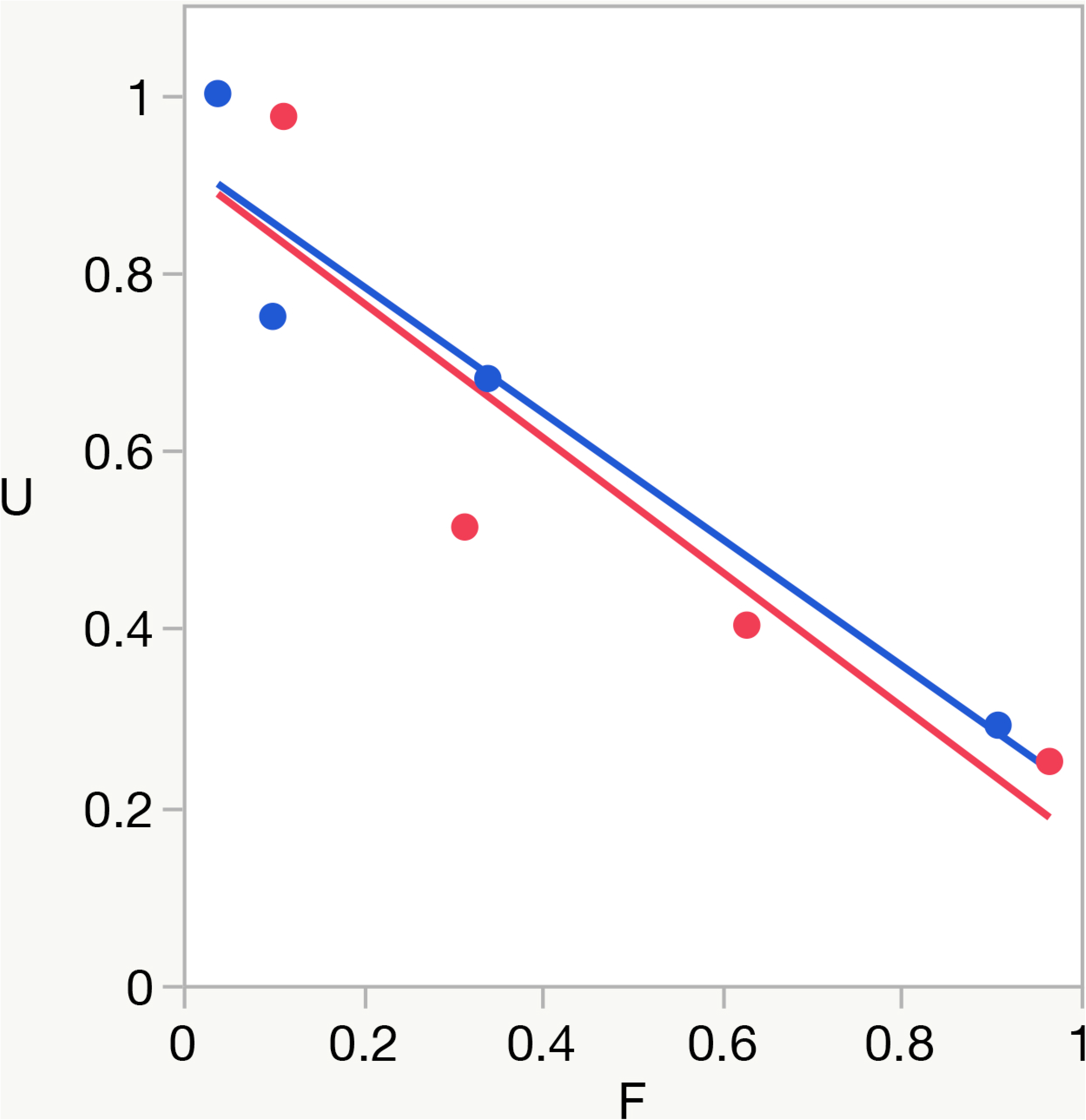
Possession of the *fli1a-F-hsp70l:Gal4VP16* transgene does not appear to increase developmental instability of the penetrance traits. Plot of proportion of unilateral expression of *fras1* mutant traits (U) as a function of trait frequency (F). Ordinary least squares regression lines for transgene-containing sets (red data points and regression lines) and groups without the transgene (blue). Slopes for both relationships are essentially identical (−0.76 and −0.71 respectively), suggesting no difference in developmental instability. The data are from the large analysis shown in Figure 7, and the different points indicate different traits as set out there. **Calculations** (from Hallgrimsson *et al*., 2005):We consider proportions, normalized such that a+b+c+d=1 for a given trait (e.g. an sy-ch fusion) and for each given condition (here, 2 conditions: mutants with included transgene compared with mutants without the transgene). Let a=neither the left nor right sides express the trait, b=only the left side expresses the trait, c=only the right side expresses the trait, d=both left and right sides express the trait. Then F= b+c+d and U= (b+c)/(b+c+d).

## DISCUSSION

### 4.1 Sequences within the *fli1a-F-hsp70l:Gal4VP16* transgene may be responsible for increased penetrance

We report here that a transgene, *fli1a-F-hsp70l:Gal4VP16*, integrated into the genome of *fras1* mutants, enhances mutant craniofacial skeletal phenotypes. This finding is highly likely to be of more general importance than applying to just this specific transgene and this specific mutation. For example, cellular stress may be involved, and the neural crest, that contributes to the cranial facial skeleton, may be particularly sensitive to cellular stress (Jones *et al*., 2008; Calo *et al*., 2018) The Gal4-UAS system is widely used in biological studies, and our finding an unexpected dominant effect on severities of phenotypes that initially were themselves the principal subject of under investigation, clearly suggests that the system, and perhaps transgenesis generally, including both in the laboratory and certainly also in the food industry, needs to be used with caution. More than this, it is possible that what we have found with a transgene introduced in the laboratory has application to a variety of transposons present normally in the genetic background. In this respect we note that the *Tol2* component of *fli1a-F-hsp70l:Gal4VP16*, responsible for its efficient transgenesis, comes from a ‘natural’ transposon, originally isolated from the medaka *Oryzias lapipes* (Kwan *et al*., 2007). Notably, half of the human genome consists of transposable elements (Lander *et al*., 2001), foreign in the same sense as is *fli1a-F-hsp70l:Gal4VP16*. Some of these elements can mobilize in nature, i.e., they are capable of hopping to new sites where they might affect host gene function, local or more globally (Munoz-Lopez & Garcia-Perez, 2010). There is a long list in the literature of causal interactions of transposons with host sequences are associated with human disease, e.g., cystic fibrosis, various cancers (Reviews; Belancio *et al*., 2008; Burns, 2017). On the other hand, transposons/transgenes might impart beneficial effects on their host (Sayah *et al*., 2004; Clark *et al*. 2006, Herpin *et al*., 2010), as could be important in evolution, but there is not much evidence for beneficial interaction. The *fli1a-F-hsp70l:Gal4VP16*–*fras1* system, with continued study, might serve as a useful model for understanding the diverse effects of such agents. In particular the remarkable diversity we encountered in the extent of the enhancements among six independent lines, all derived from the same zebrafish strain injected with the same plasmid, but with different insertion sites, bears further examination. The fact that we observed increased penetrance in mutants bearing independent transgene integrations makes it extremely unlikely that disruption of host sequences adjacent to or near the insertion sites are responsible for the enhancement. Rather, our evidence points to sequences within the transgene itself being causal.

As pointed out in the Introduction, precedents make it reasonable to suggest that sequences within the *fli1a-F-hsp70l:Gal4VP16* itself are responsible for the observed penetrance enhancements. Namely, the known action of squelching of transcription by Gal4VP16 (Gill & Ptashne, 1988), suggested because it is an off-target activity, could somehow potentiate the loss of function of *fras1*. It is particularly mysterious, and of considerably interest that the enhancements are specific, in the sense that we don’t see new phenotypes (e.g. a fusion of the joint in the second arch) that are not present in wild types or in *fras1* homozygous mutants. Two of the traits studied here (sy-pq fusion, op shape) occur at very low frequency in the *fras1* mutants in the absence of the transgene, but they are present in mutants nevertheless, and enhanced by the transgene. We found no evidence of toxicity, developmental delay, or other features that may suggest generalized abnormalities, and we note that Acridine Orange staining suggested that apoptosis is low in the pharyngeal arches, with no discernable difference due to the transgene.

### 4.2 In what tissue might the transgene act to enhance *fras1* mutant skeletal phenotypes?

To understand how the transgene might be acting, we need to consider what specific developmental process or processes might the transgene affect that could lead to increased penetrance of the mutant phenotypes? Transgene induced variation of outpocketing of the late-forming region of the first pharyngeal pouch (late p1) is unlikely to be responsible for enhancement since loss of outpocketing is fully penetrant in *fras1* mutants without the transgene being present (Talbot *et al*., 2012). Further arguing against an endodermal site for the action of the transgene, we note that *fli1a* is expressed in pharyngeal arch mesenchyme, not endoderm; hence we do not expect the *fli1a-F-hsp70l:Gal4VP16* to be active (and exert its enhancing effect) in endoderm.

Instead of the endoderm, the transgene could be exerting its effects directly in the (mesenchymal) skeletal forming cells themselves. However, it seems unlikely that the mechanism is the same for all of the enhanced phenotypes we described. The sy-ch cartilage fusion occurs at an ectopic location between cartilages that are not normally close neighbors but become so in *fras1* mutants, in part due the loss of late p1 which normally separates them. In contrast, the M-pq fusion is at the location where the two cartilages normally articulate – forming the first arch (jaw) joint. The joint normally is carved out of a single precartilage condensation that includes precursors of both the M and pq cartilages. Joint formation is under control of endodermal Edn1-dependent transcription factors including Hand2 (positively regulating joint formation) and Barx1 (negatively regulating; Nichols *et al*., 2013). It is conceivable that failure of endodermal late p1 outpocketing provides an environment conducive for the enhancements of the two fusions (sy-ch and M-pq) for completely different reasons: change in spatial relationships between cartilages on the one hand (sy-ch), and change in Edn1 signaling from the endoderm on the other (M-pq). If this were true, then the molecular basis of enhancement by the transgene could be quite different for the two fusions. We note that the second arch joint, involving the hyosymplectic, interhyal, and ceratohyal, and developing only a few cell diameters away from the site of the *fras1*^−^ sy-ch fusion, is normal in *fras1* mutants, including mutants possessing the *fli1a-F-hsp70l:Gal4VP16* transgene. In spite of this small distance it could be that endodermal signaling to the second arch joint-forming cells is from second arch endoderm, not late p1, possibly explaining why this joint region does not exhibit the fusion. The *fras1* mutant phenotypes, as well as their enhancement by the transgene are exquisitely specific.

The strong enhancement of the op shape mutant phenotype in our original line A, and in two of the new lines (B, E; Figure 7) likely has nothing to do with late p1 or endoderm at all. The op is a dermal bone, the designation as dermal signifying its origin in the mesenchyme of the dermis, closely adjacent to second arch ectoderm, but not particularly close to endoderm. Ectoderm (like endoderm) expresses *fras1*, and its loss from an ectodermal source in *fras1* mutants is apparently responsible for the fin defects which first led to the identification of the mutant in zebrafish (Carney et al., 2010). The defect in op morphology is encountered only rarely in the absence of the transgene. Among all lines studied here, and including both left and right sides, for the dataset used for Figure 7, the stick morphology was present in only 44/555 left or right sides (7.9%) when the transgene was absent, and only 1/238 sides of phenotypically wild-type larvae (i.e., larvae homozygous or heterozygous for the *fras1*^+^ allele, 0.4%). The rarity likely explains why the phenotype was not discovered in our previous studies (Talbot *et al*., 2012), and only part way through the current study. The low frequency shows that the transgene can apparently effect skeletal cells in spite of development occurring mostly normally in mutants. In summary, we think it likely that enhancement of mutant phenotypes is occurring within the skeletogenetic mesenchyme, but probably by different mechanisms for the different phenotypes.

### 4.3 Enhancement could be due to retardation or truncation of development

The op could provide clues as to the nature of the enhancement. In the severe enhanced mutants, a ‘stick’ shape replaces the fan shape normally present at this stage. We consider two hypotheses. One, in which developmental timing is important is that in normal development a stick shape appears before the fan shape. The stick shape persists for about a day, the bone elongating during this time, before a new pattern of osteoblast activity appears and the bone begins to transform to the fan shape (Kimmel *et al*., 2010). Hence, we can propose that a retardation or truncation of development by the transgene prevents the stick-to-fan switch, and, in effect, is the basis of the enhancement. We can shoehorn one of the cartilage fusions into the same retardation/truncation hypothesis – the M-pq begins development as a single pre-cartilage element, that likely persists as the ancestral state, because in the lamprey (an agnathan) the first arch cartilage is not jointed (Cerney *et al*., 2010, Nichols *et al*., 2013). In gnathostomes, including zebrafish in particular, the joint region is carved out secondarily as a new development. Loss of this later phase in *fras1* mutants, the loss enhanced by the transgene, would explain the observed phenotype.

The retardation/truncation hypothesis does not perfectly explain the data, however. For one thing, the genes regulating the the first arch joint formation and the op shape transition, and their timing, appear to differ for the op (e.g., local upregulation of hedgehog signaling; Huycke *et al*., 2012) and the M-pq (local downregulation of *barx1*, Nichols *et al*., 2013). Furthermore, the hypothesis does not readily explain the very frequent enhancement of the sy-ch fusion as well as the much rarer sy-pq fusion. For at least the sy-ch fusion, penetrance increases during development in *fras1* mutants (Talbot *et al*., 2012), whereas retardation/truncation predicts decrease.

### 4.4 Enhancement could be due to dorsal-ventral (DV) homeosis

A second hypothesis coming from op shape is based on homeosis. The op is transformed from a fan to a stick shape in mutants with loss of function of genes of the jagged-Notch pathway, and considerable evidence is available that this pathway functions in specifying dorsal skeletal identity (Zuniga *et al*., 2010; 2011; Barske *et al*., 2016). The transformation thus is toward a more ventral element, a branchiostegal ray, which is stick-shaped in the wild type. For example, a branchiostegal ray is Alizarin Red stained in Figure 1 (br). With respect to the cartilage phenotypes, skeletal fusions often accompany homeosis, e.g. fusion of adjacent vertebrae and limb skeletal elements in Hox mutants (review: Quinonez & Innis, 2014). Notably, DV homeotic mutations, e.g., loss of Dlx5 and Dlx6 in the mouse, can result in fusion of putative more dorsal and more ventral first pharyngeal arch elements, the incus and malleus (Depew *et al*., 2002). Similarly, in zebrafish, loss of Dlx gene function (Talbot *et al*., 2010) as well as *edn1* mutation (Miller *et al*., 2000; Walker *et al*., 2006) can result in fusion of dorsal and ventral elements homologous to the mammalian malleus and incus in the first arch (M-pq fusion, as we also see in *fras1* mutants) and, as well, fusion of the DV joint in second arch. The sy-ch fusion of the second arch in *fras1* mutants also is between elements with different DV identities, sy being more dorsal than the ch (Schilling & Kimmel, 1997). These features make the DV homeosis hypothesis for the transgene enhanced disturbance in *fras1* mutants an attractive one. As for the retardation/truncation hypothesis, the homeosis hypothesis can be tested by developmental genetics – in particular, by examining homeotic patterning gene expression in the *fras1* mutants, and by looking genetically at the interaction of *fras1* with patterning genes. A difficulty is that except for the op, we do not see skeletal shape changes characteristic of homeotic mutants in the *fras1* mutants, with or without the transgene. The transformations would seemingly be very local, in the neighborhood where late p1 would be located.

### 4.5 Loss of canalization explains enhancement variation and specificity

The phenotypes of normal (non-mutant) individuals exhibit developmental robustness – resistance to perturbations that could arise from mutations in the genetic background or from environmental insults. In insightful analyses conducted during the middle of the last century, C. Waddington showed that whereas phenotypes are robust in wild-type organisms (fruit flies in his case), robustness, which he termed “canalization” can be strikingly reduced in mutants, and also in severe environmental disturbances, such that phenotypic variation is increased (Waddington, 1942; 1957). Such loss of canalization, as we might take to be the case in *fras1* mutants, occurs for phenotypes specific to the de-canalizing mutation (*fras1* in our case) not generally across the tissues of the organism. Waddington imagined that robustness of a particular wild-type phenotype results from the output of a specific network of genes, evolved under the influence what we would now call stabilizing selection, and that deleterious mutations can unbalance the network. Canalization has had many proponents since Waddington’s day (Hallgrimsson *et al*., 2002; 2006; Dworkin, 2005; Flatt, 2005), but its mechanism is still poorly understood. The epigenome has provided a recent alternative (or additional) attractive explanation for the phenomenon (Pujadas & Feinberg, 2012). Our result suggesting that developmental instability is unchanged by the transgene (Figure 8) does not necessarily argue against canalization being involved: Waddington thought that buffering of developmental noise was by a different mechanism than canalization – buffering the effects of a genetic or environmental disturbance (Waddington, 1957; 1960; Waddington & Robertson, 1966). However, whether canalization and the buffering of developmental noise are genetically the same or different is currently controversial (e.g., Debat *et al*., 2000; 2009; Hallgrimsson *et al*., 2002; Dworkin, 2005; Breuker *et al*., 2006). Loss of canalization provides a relevant hypothesis to account especially for the specificity of the enhancing action of the transgene. The hypothesis is not mutually exclusive with the two described above.

To illustrate the canalization-loss hypothesis, we imagine a scenario where squelching is the root cause of enhancement and varies according to the level of transgene expression. We would expect squelching, at the transcriptional level, in those cells expressing the transgene at high levels, as in craniofacial skeleton-forming cells. The most sensitive phenotypes would be those for which canalization is decreased, as the hypothesis proposes for Fras1-dependent functions in *fras1* mutants. For phenotypes not dependent on Fras1, resistant to squelching or its effects would be observed because these phenotypes would still be well buffered. Hence the hypothesis accounts for the specificity we observe of the enhancements in transgene-bearing mutants. The hypothesis predicts phenotypes to be more severe with higher transgene dosage, as our incross-outcross comparison indicates. The canalization-loss model also can explain why the phenotypic enhancements vary among our transgenic lines; we need only to assume that, for whatever reason (including position effects imparted by the insertion site), the level of transgene function varies between the lines, as would underlie the severity differences.

### 4.6 Conclusion

We show here that *fras1* mutant phenotypes are sensitive to the presence of an integrated transgene, *fli1a-F-hsp70l:Gal4VP16*, in the genetic background. The effect is apparently due to an off-target transgene function itself (such as squelching), rather than to change in function of host sequences near the insertion site of the transgene. Specific *fras1* mutant phenotypes are made more severe, in the apparent absence of other disruptions, including malformation and developmental retardation, that would indicate toxicity. We hypothesize that loss of canalization explains the variation among our transgenic lines, and accounts for the specificity. Furthermore, along with known issues concerning off-target effects of the transgene, loss of canalization points to misregulated transcription as being the proximate cause of the phenotypic enhancements. Because of the low background of generalized effects, this system would seemingly be a good one for transcriptomic studies to identify the genes important in the regulation of skeletal phenotypic variation.

## ACKNOWLEDGEMENTS

We thank members of the Institute of Neuroscience, Univ. of Oregon for discussion and comment. D. Farnsworth commented on an early version of the manuscript. Staff of the Univ. of Oregon Aquatic Animal Care Services provided excellent animal care and husbandry. G. Crump and M. Matsutani provided transgenic fish and plasmids. We are grateful to D. Turnbull and M. Weitzman for sequencing.

## AUTHOR CONTRIBUTIONS

C.B.K.: designed the experiments. C.B.K., J.H.P. and J.T.N contributed ideas and to manuscript preparation. W.O., S.A., C.W., J.D., B.B.-S., T.A.T. and J.T.N. generated the data. C.B.K., A.L.W., P.B. C.W., J.D. and J.T.N. contributed to data analysis. C.B.K. and J.H.P. obtained funding. All authors reviewed and approved the manuscript.

## SIGNIFICANCE STATEMENT

Transgenes are widespread in animal genomes, and transgenesis is widespread in biological research, and in the food industry; yet transgenes are frequently ignored in scientific work. To provide a model for understanding transgene-host interaction we have demonstrated increased severity of zebrafish *fras1* mutant skeletal phenotypes in the presence of a potent Gal4 transgene known to effect transcription. We hypothesize that the phenotypes are enhanced by the transgene due to the loss of genetic robustness.

## REFERENCES

Asakawa, K., & Kawakami, K. (2009). The Tol2-mediated Gal4-UAS method for gene and enhancer trapping in zebrafish. Methods, 49, 275–281

Asakawa, K., Suster, M. L., Mizusawa, K., Nagayoshi, S., Kotani, T., Urasaki, A., Kishimoto, Y., Hibi,M., & Kawakami, K. (2008). Genetic dissection of neural circuits by Tol2 transposon-mediated Gal4 gene and enhancer trapping in zebrafish. Proceedings of the National Academy of Sciences of the United States of America, 105, 1255–1260

Barske, L., Askary A., Zuniga, E., Balczerski, B., Bump, P., Nichols, J.T., Crump, J.G. (2016). Competition between Jagged-Notch and Endothelin1 signaling selectively restricts cartilage formation in the zebrafish upper face. PLoS Genetics, 12, e1005967

Belancio, V.P., Hedges, D.J., & Deininger, P. (2008). The Tol2-mediated Gal4-UAS method for gene and enhancer trapping in zebrafish. Genome Research, 18, 343–358

Bolger, A.M., M. Lohse, M., & Usadel, B. (2014). Trimmomatic: a flexible trimmer for Illumina sequence data. Bioinformatics, 30, 2114–2120

Breuker, C.J., Patterson, J.S., Klingenberg, C.P. (2006). A single basis for developmental buffering of *Drosophila* wing shape, PLoS One, 1, e7

Burns, K. H. (2017) Transposable elements in cancer. Nature Reviews Cancer, 17, 415–424

Calo, E., Gu, B., Bowen, M.E., Aryan, F., Zalc, A., Liang, J., Flynn, R.A., Swigut, T., Chang, H.Y., Attardi, L.D., & Wysocka, J. (2018) Tissue-selective effects of nucleolar stress and rDNA damage in developmental disorders. Nature, 554, 112–117

Carney, T.J., Feitosa, N.M., Sonntag, C., Slanchev, K., Kluger, J., Kiyozumi, D., Gebauer, J.M., Talbot, J.C., Kimmel, C.B., Sekiguchi, K., Wagener, R., Schwarz, H., Ingham, P.W., & and Hammerschmidt. M. (2010). Genetic analysis of fin development in zebrafish identifies furin and hemicentin1 as potential novel fraser syndrome disease genes. PLoS Genetics, 6, e1000907

Cerny, R., Cattell, M., Sauka-Spengler, T., Bronner-Fraser, M., Yu, F. Q. and Medeiros, D. M. (2010). Evidence for the prepattern/cooption model of vertebrate jaw evolution. Proceedings of the National Academy of Sciences of the United States of America, 107, 17262–17267

Clark, L.A., Wahl, J.M., Rees, C.A., & Murphy, K.E. (2006). Retrotransposon insertion in SILV is responsible for merle patterning of the domestic dog. Proceedings of the National Academy of Sciences of the United States of America, 103, 1376–1381

Das, A., & Crump, J.G. (2012). Bmps and Id2a act upstream of Twist1 to restrict ectomesenchyme potential of the cranial neural crest. PLoS Genetics, 8, e1002710

Debat, V., Alibert, P., Paradis, E., Auffray, J.-C. (2000). Independence between developmental stability and canalization in the skull of the house mouse. Proceedings of the Royal Society B Biological Sciences, 267, 423–430

Debat, V., Debelle, A., Dworkin, I. (2009). Plasticity, canalization, and developmental stability of the *Drosophila* wing: joint effects of mutations and developmental temperature. Evolution, 63, 2864–2876.

Depew, M.J., Lufkin, T., Rubenstein, J.L.R. (2002). Specification of jaw subdivisions by Dlx genes. Science, 298, 381–384

Dongen, S. V. (2006). Fluctuating asymmetry and developmental instability in evolutionary biology: past, present and future. Journal of Evolutionary Biology, 19, 1727–1743

Dworkin, I. (2005). Canalization cryptic variation, and developmental buffering: a critical examination and analytical perspective. In B. Hallgrímsson, B. K. Hall (Eds.), Variation: A central concept in biology (pp. 131–158). Amsterdam, Netherlands, Elsevier Academic Press

Flatt, T. (2005). The evolutionary genetics of canalization. The Quarterly Review of Biology, 80, 287–316

Ghosh, A., & Halpern, M.E. (2016). Transcriptional regulation using the Q system in transgenic zebrafish. Methods in Cell Biology, 135, 205–218

Gill, M., & Ptashne, M. (1988). Negative effect of the transcriptional activator GAL4. Nature, 334, 721–724

Graham, J.H., Raz, S., Hel-Or, H., & Nevo, E. (2010). Fluctuating asymmetry: methods, theory, and applications. Symmetry, 2, 466–540

Hallgrımsson, B., Brown, J.J.Y., Ford-Hutchinson, A.F., Sheets, H.D., Zelditch, M.L., and & Jirik, F.R. (2006). The brachymorph mouse and the developmental-genetic basis for canalization and morphological integration. Evolution and Development, 8, 61–73

Hallgrimsson, B., & Hall, B.K., eds. (2005) Variation: A central concept in biology. Amsterdam, Netherlands, Elsevier Academic Press

Hallgrimsson, B., O’Donnabhain, B., Blom, D.E., Willmore, K.E. (2005). Why are rare traits unilaterally expressed? Trait frequency and unilateral expression for cranial nonmetric traits in humans. American Journal of Physical Anthropology, 128, 14–25

Hallgrimsson, B., Willmore, K., Hall, B.K. (2002). Canalization, developmental stability, and morphological integration in primate limbs. American Journal of Physical Anthropology, 119, 131–158

Herpin, A., Braasch, I., Kraeussling, M., Schmidt, C., Thoma, E.C.,Nakamura, S., Tanaka, M., & Schartl, M. (2010). Transcriptional rewiring of the sex determining *dmrt1* gene duplicate by transposable elements. PLoS Genetics, 6, e1000844

Howe, K., et al. (2013). The zebrafish reference genome sequence and its relationship to the human genome. Nature, 496, 498–503

Huycke, T.R., Eames, B.F., & Kimmel, C.B. (2012). Hedgehog-dependent proliferation drives modular growth during morphogenesis of a dermal bone Development, 139, 2371–2380

Jones, N.C., Lynn, M. L., Gaudenz, K., Sakai, D., Aoto, K., Rey, J.-P., Glynn, E.F., Ellington, L., Du, C., Dixon, J., Dixon, M.J., & Trainor, P.A. (2008) Prevention of the neurocristopathy Treacher Collins syndrome through inhibition of p53 function. Nature Medicine, 14, 125–133

Kimmel, C.B., DeLaurier, A., Ullmann, B., Dowd, J., & McFadden, M. (2010). Modes of developmental outgrowth and shaping of a craniofacial bone in zebrafish. PLoS One, 5, e9475

Koster, R.W., & Fraser, S.E. (2001). Tracing transgene expression in living zebrafish embryos. Developmental Biology, 233, 329–346

Kwan, K.M., Fujimoto, E., Grabher, C., Mangum, B.D., Hardy, M.E., Campbell, D.S., Parant, J.M., Yost, H.J., Kanki, J.P., and Chien, C.-B. (2007) The Tol2kit: a multisite gateway-based construction kit for Tol2 transposon transgenesis constructs. Developmental Dynamics, 236, 3088–3099

Lander, E.S., Linton, L.M., Birren, B., Nusbaum, C., Zody, M.C., Baldwin, J., Devon, K., Dewar, K., Doyle, M., FitzHugh, W., et al. (2001). Initial sequencing and analysis of the human genome. Nature, 409, 860–921

Meyer, S.N., Amoyel, M., Bergantinos, C., de la Cova, C., Schertl, K., Basler, K., & Johnston, L.A. (2014). An ancient defense system eliminates unfit cells from developing tissues during cell competition. Science, 346, (6214), 1258236-1–1258236-8

Miller, C.T., Schilling, T.F., Lee, K.-H., Parker J., & Kimmel, C.B. (2000). *sucker* encodes a zebrafish Endothelin-1 required for ventral pharyngeal arch development. Development, 127, 3815–3828

Munoz-Lopez M., & Garcia-Perez, J. L. (2010). DNA Transposons: Nature and Applications in Genomics. Current Genomics, 11, 115–128

Nichols, J.T., Pan L., Moens, C.B., & Kimmel, C.B. (2013). *barx1* represses joints and promotes cartilage in the craniofacial skeleton. Development, 140, 2765–2775

Polak, M. (Ed.) (2003). Developmental instability: causes and consequences. Oxford, UK: Oxford University Press

Pujadas, E., & Feinberg, A.P. (2012). Regulated noise in the epigenetic landscape of development and disease. Cell, 148, 1123–1131

Quinonez, S.C., & Innis, J.W. (2014). Human HOX gene disorders. Molecular Genetics and Metabolism, 111, 4–15

Sadowski, I., Ma, J., Triezenberg, S., & Ptashne, M. (1988). GAL4-VP16 is an unusually potent transcriptional activator. Nature, 335, 563–564

Sayah, D.M., Sokolskaja, E., Berthoux, L., and Luban, J. (2004). Cyclophilin A retrotransposition into TRIM5 explains owl monkey resistance to HIV-1. Nature, 430, 569–573

Schilling T.F., & Kimmel, C.B. (1997). Musculoskeletal patterning in the pharyngeal segments of the zebrafish embryo. Development, 124, 2945–2960

Scott, E.K., Mason, L., Arrenberg, A.B., Ziv, L., Gosse, N.J., Xiao, T., Chi, N.C., Asaka, K., Kawakami, K., & Baier, H. (2007). Targeting neural circuitry in zebrafish using GAL4 enhancer trapping. Nature Methods, 4, 323–326

Slavotinek, A., Li, C., Sherr, E. H. and Chudley, A. E. (2006). Mutation analysis of the FRAS1 gene demonstrates new mutations in a propositus with Fraser syndrome. American Journal of Medical Genetics A, 140, 1909–1914

Smyth, I. and Scambler, P. (2005). The genetics of Fraser syndrome and the blebs mouse mutants. Human Molecular Genetics, 14, R269–R274

Talbot J.C., Johnson, S.L., & Kimmel, C.B. (2010). *hand2* and *Dlx* genes specify dorsal, intermediate and ventral domains within zebrafish pharyngeal arches. Development, 137, 2507–2517

Talbot, J.C., Nichols, J.T., Yan Y.-L., Leonard, I.F., BreMiller, R.A., Amacher, S.L., Postlethwait, J.H., Kimmel, C.B. (2016). Pharyngeal morphogenesis requires *fras1-itga8*-dependent epithelial-mesenchymal interaction. Developmental Biology, 416, 136–148

Talbot, J.C., Walker, M.B., Carney, T.J., Huycke, T.R., Yan, Y.-L., Bremiller, R.A., Gai, L., Delaurier, A., Postlethwait, J.H., Hammerschmidt, M., & Kimmel, C.B. (2012). *fras1* shapes endodermal pouch 1 and stabilizes zebrafish pharyngeal skeletal development. Development, 139, 2804–2813

Tucker, B., & Lardelli, M. (2007). A rapid apoptosis assay measuring relative Acridine Orange fluorescence in zebrafish embryos. Zebrafish, 4, 113–116

Waddington, C.H. (1942). Canalization of development and the inheritance of acquired characters. Nature, 150, 563–565

Waddington, C.H. (1957). The Strategy of the genes: A discussion of some aspects of theoretical biology. London, UK. Allen & Unwin

Waddington, C.H. (1960). Experiments on canalizing selection. Genetics Research (Cambridge), 1, 140–150

Waddington, C.H. & Robertson, E. (1966). Selection for developmental canalization. Genetics Research (Cambridge), 7, 303–312

Walker, M.B., & Kimmel, C.B. (2007). A two-color acid-free cartilage and bone stain for zebrafish larvae. Biotechnic & Histochemistry, 82, 23–28

Walker, M.B., Miller, C.T., Talbot, J.C., Stock, D.W., & Kimmel, C.B. (2006). Zebrafish *furin* mutants reveal intricacies in regulating Endothelin1 signaling in craniofacial patterning. Developmental Biology, 295, 194–205

Westerfield, M. (2007). The zebrafish book. A guide for the laboratory use of zebrafish (*Danio rerio*). 4th ed., Eugene, OR. Univ. of Oregon Press

Wu, T.D., Reeder, J., Lawrence, M., Becker, G. (2016). GMAP and GSNAP for genomic sequence alignment: Enhancements to speed, accuracy, and functionality. Methods in Molecular Biology, 1418, 283–334

Zuniga, E., Rippen, M., Alexander, C., Schilling, T.F., & Crump, J.G. (2011). Gremlin 2 regulates distinct roles of BMP and Endothelin 1 signaling in dorsoventral patterning of the facial skeleton. Development, 138, 5147–5156

Zuniga, E., Stellabotte, F., & Crump, J.G. (2010). Jagged-Notch signaling ensures dorsal skeletal identity in the vertebrate face. Development, 137, 1843–1852

